# Causal considerations can determine the utility of machine learning assisted GWAS

**DOI:** 10.1101/2024.12.16.628604

**Authors:** Sumit Mukherjee, Zachary McCaw, David Amar, Rounak Dey, Thomas Soare, Kaiwen Xu, Hari Somineni, insitro Research Team, Nicholas Eriksson, Colm O’Dushlaine

**Affiliations:** insitro inc., South San Francisco, California, USA

**Keywords:** Machine Learning derived phenotype, GWAS, Imputation, Prediction-based Inference

## Abstract

Machine Learning (ML) is increasingly employed to generate phenotypes for genetic discovery, either by imputing existing phenotypes into larger cohorts or by creating novel phenotypes. While these ML-derived phenotypes can significantly increase sample size, and thereby empower genetic discovery, they can also inflate the false discovery rate (FDR). Recent research has focused on developing estimators that leverage both true and machine-learned phenotypes to properly control the type-I error. Our work complements these efforts by exploring how the true positive rate (TPR) and FDR depend on the causal relationships among the inputs to the ML model, the true phenotypes, and the environment.

Using a simulation-based framework, we study architectures in which the machine-learned proxy phenotype is derived from biomarkers (i.e. inputs) either causally upstream or downstream of the target phenotype. We show that no inflation of the false discovery rate occurs when the proxy phenotype is generated from upstream biomarkers, but that false discoveries can occur when the proxy phenotype is generated from downstream biomarkers. Next, we show that power to detect variants truly associated with the target phenotype depends on its heritability and correlation with the proxy phenotype. However, the source of the correlation is key to evaluating a proxy phenotype’s utility for genetic discovery. We demonstrate that evaluating machine-learned proxy phenotypes using out-of-sample predictive performance (e.g. phenotypic correlation) provides a poor lens on utility. This is because overall predictive performance does not differentiate between genetic and environmental correlation. In addition to parsing these properties of machine-learned phenotypes via simulations, we further illustrate them using real-world data from the UK Biobank.

## 1 Introduction

Genome-wide association studies (GWAS) identify genetic variations that are associated with a particular phenotype, and have revolutionized our understanding of the genetic architecture of complex traits and diseases [1]. This approach has successfully uncovered numerous genetic variants that contribute to the risk of complex disorders, paving the way for precision medicine and targeted therapeutic interventions. The advent of large-scale biobanks has further propelled the field of genetic discovery by providing structured, well-powered, data sets with genotypes and deep phenotyping for hundreds of thousands of individuals [2, 3, 4, 5]. The quality and unprecedented scale of these data sets have empowered researchers to detect genetic variants with ever smaller effect sizes while capturing an ever greater proportion of complex trait heritability [6].

Despite the size and scope of population-based biobanks, the challenge of data sparsity persists. Many phenotypes of interest are measured for only a subset of participants, and only a fraction of participants will develop any given disease, limiting the effective sample size available for GWAS in observational biobanks. This sparsity can meaningfully diminish the power to detect associations for complex traits with modest genetic effects. To address this limitation, researchers increasingly turn to machine learning (ML) to impute missing phenotypic values from the available data. These ML imputation approaches have been shown to improve genetic discovery for difficult to ascertain phenotypes, such as the optical cup-to-disc ratio, thoracic aortic diameter, major depression, and hepatic fat percentage [7, 8, 9, 10, 11, 12, 10].

When performing GWAS on predicted or imputed outcomes, the relationship between genotype and the imputed phenotype may differ both quantitatively and qualitatively (i.e. in existence or direction) from that between genotype and the target outcome [13, 14, 15]. Distortion of the genotype-phenotype relationship due to imputation can lead to inflated type I error, due to the detection of signals that spuriously associate with the imputed phenotype but not the true phenotype, and compromise the utility of downstream analyses that depend on unbiased effect size estimation, such as polygenic scoring. Recent work [16, 17, 13, 14, 15] has focused on developing methods to address these challenges and minimize the risk of false discoveries. Specifically, these methods aim to provide unbiased estimation for the effect of genotype on the target outcome that is robust to the accuracy or quality of the imputation model. In doing so, these methods guarantee valid inference, but may forego power as compared with the simpler strategy of proxy GWAS (**Figure 1**), where the machine-learned proxy phenotype is studied in place of the original phenotype.

**Figure 1:**
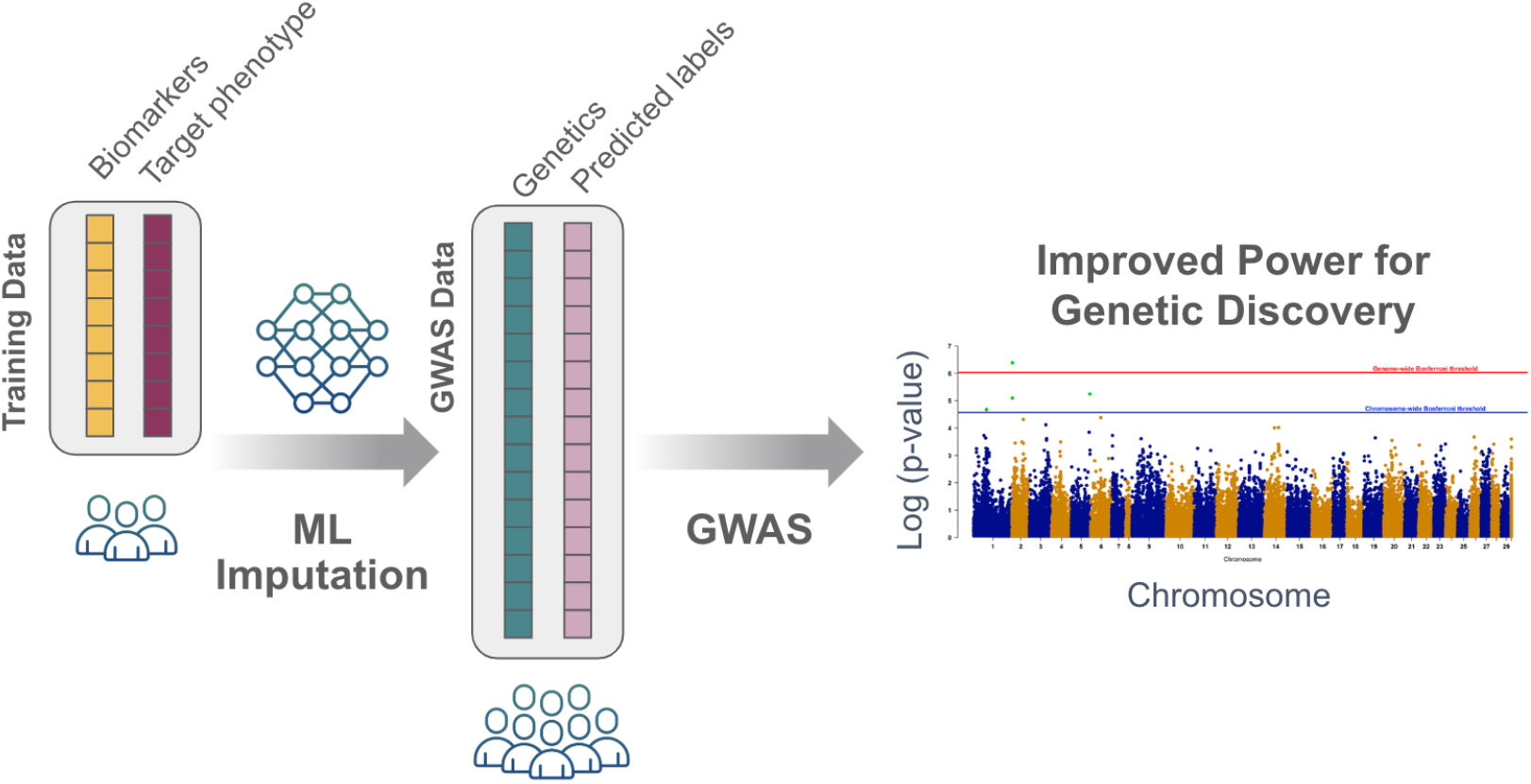
Typical setup for machine learning (ML)-assisted proxy GWAS. Within a labeled training data set, surrogate biometric data (e.g. biomarkers) is utilized to develop models for imputing the target phenotype. Once trained, the model extrapolates phenotypic labels across a larger dataset, augmenting the effective sample size for genome-wide association studies (GWAS). Since the predicted labels are used in place of measured, ground-truth labels, this strategy is referred to as proxy GWAS. The goal is to leverage the imputed proxy phenotype to identify genetic variants associated with the original target phenotype.

This work is intended to complement the recent work in prediction-based inference by studying how the power and false discovery rate of proxy GWAS depend on the causal relationships among genotypes, the true phenotype, and the variables the enter the imputation model, which we describe as biomarkers. The remainder of this paper is organized as follows. Section 2 describes our simulation framework and details of the real data analysis. Section 3 includes our case-studies on simulated and real data. We first contrast proxy phenotypes imputed from upstream versus downstream biomarkers in terms of their power for detecting true positive association and their FDR. Second, we examine how heritability of the true phenotype and its correlation with the proxy phenotype affect power to recover true positive associations. Third, we decompose the phenotypic correlation, assessing the dependence of the true positive rate (TPR) and the FDR on genetic versus environmental correlations. We conclude with the implications of our case-studies for practice.

## 2 Data and Methods

### 2.1 Real-world dataset

The UK Biobank (UKB) is a large-scale prospective cohort study and biomedical database containing detailed genetic and health information on approximately 500,000 individuals from the United Kingdom, aged between 40 and 69 years at the time of recruitment [18]. The UKB resource encompasses a wide variety of data on health-related outcomes, including hospital records, cancer registries, death records, and physical measurements, as well as selfreported health questionnaires. Additionally, the resource contains a vast array of biological measurements such as blood, urine, and saliva biochemistry.

### 2.2 Methods for simulated traits

#### 2.2.1 Simulating traits with a specified heritability and genetic correlation

Let **G** denote a vector of *J* genetic variants in linkage equilibrium, with elements *G*_*j*_ *∼* Binom(2, *p*), where *p* is the minor allele frequency. ***G*** can contain both causal and noncausal variants for biomarkers and phenotypes. Let *S*_bio_ and *S*_pheno_ denote the indices of the causal variants for biomarkers and phenotypes. Non-causal variants have effect sizes of zero and are therefore not included in either *S*_bio_ or *S*_pheno_. The sets *S*_bio_ and *S*_pheno_ may or may not overlap.

For the *j*th variant causal for biomarkers, let ***β***_bio,*j*_ denote an *n*_bio_ *×* 1 vector representing the non-zero effects of the *j*th variant on the *n*_bio_ biomarkers. Similarly, let ***β***_pheno,*j*_ denote the *n*_pheno_ *×*1 vector of non-zero effects for the *j*th variant in *S*_pheno_ on the *n*_pheno_ phenotypes. We assume that for each causal variant, the vectors ***β***_bio,*j*_ and ***β***_pheno,*j*_ follow a joint multivariate normal distribution:

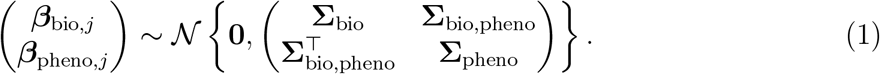

Here, **Σ**_bio_ is an *n*_bio_ *× n*_bio_ diagonal matrix with the variances of the effect sizes for each biomarker on the diagonal. The diagonal elements 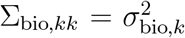 determine the contribu-tion of the *j*th genetic variant to the heritability of the *k*th biomarker. Likewise, **Σ**_pheno_ is a *n*_pheno_ *× n*_pheno_ diagonal matrix with the effect size variances for each phenotype on the diagonal. The diagonal elements 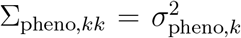 determine the contribution of the genetic variance to the heritability of the *k*th phenotype. Lastly, **Σ**_bio,pheno_ is the *n*_bio_ *× n*_pheno_ matrix where each element Σ_bio,pheno,*kk*_*′* represents the covariance between the effect sizes of the *k*th biomarker and the *k*^*′*^th phenotype due to the *j*th causal variant.

The final *n*_bio_ *×* 1 biomarker vector and *n*_pheno_ *×* 1 phenotype vector are generated as:

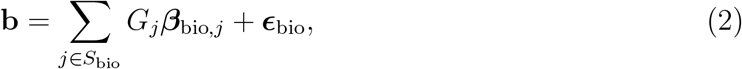

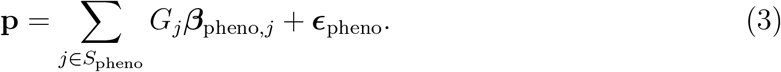

The environmental components are simulated as:

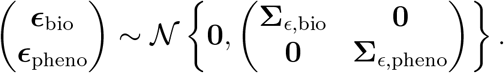

Here **Σ**_*ϵ*,bio_ and **Σ**_*ϵ*,pheno_ are diagonal *n*_bio_ *× n*_bio_ and *n*_pheno_ *× n*_pheno_ matrices defining the environmental contributions to the variances of biomarkers and phenotypes respectively.

#### 2.2.2 Generating downstream traits with specified degree of environmental influence and direct genetic effects

When generating downstream phenotypes ***p***_down_, we first simulate upstream biomarkers ***b***_up_ following (2), then construct the phenotypes as follows:

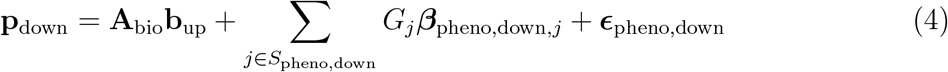

Here ***A***_bio_ is an *n*_pheno_ *× n*_bio_ matrix representing a linear relationship between upstream biomarkers and downstream phenotypes, *S*_pheno,down_ is the set of indices for the variants in ***G*** with direct causal effects on the downstream phenotype, ***β***_pheno,down,*j*_ is an *n*_pheno_ *×* 1 vector of random effect sizes drawn from a *𝒩* (**0, Σ**_pheno,down_) distribution, where **Σ**_pheno,down_ is a diagonal matrix, and ***ϵ***_pheno,down_ is an *n*_pheno_ *×* 1 residual drawn from a *𝒩* (**0, Σ**_*ϵ*,down_) distribution, where **Σ**_*ϵ*,down_ is again a diagonal matrix. The diagonal elements of **Σ**_pheno,down_ determine the contribution of direct genetic effects to the variance of ***p***_down_, while those of **Σ**_*ϵ*,down_ determine the contribution of environmental effects.

Analogously, to generate downstream biomarkers ***b***_down_, we first simulate upstream phenotypes ***p***_up_ following (3), then construct the biomarkers as follows:

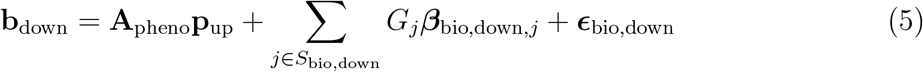

Here ***A***_pheno_ is an *n*_bio_ *× n*_pheno_ matrix representing a linear relationship between upstream phenotypes and downstream biomarkers, *S*_bio,down_ is the set of indices for the variants in ***G*** with direct causal effects on the downstream biomarkers, ***β***_bio,down,*j*_ is an *n*_bio_ *×* 1 vector of random effect sizes drawn from a 𝒩 (**0, Σ**_bio,down_) distribution, and ***ϵ***_bio,down_ is an *n*_bio_ *×* 1 residual drawn from a 𝒩 (**0, Σ**_*ϵ*,down_) distribution.

#### 2.2.3 Heritability and Genetic Correlation Estimation

The heritability for each biomarker and phenotype was estimated as the proportion of the phenotypic variance explained by the genetic variants. For a set of *N* individuals with *J* genetic variants, the genotype matrix is denoted by **G** *∈* ℝ^*N×J*^. Let 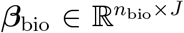 and 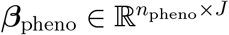 be the matrices of genetic effects for *n*_bio_ biomarkers and *n*_pheno_ phenotypes respectively. The heritability of biomarkers 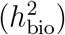 and phenotypes 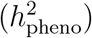 are calculated as follows:

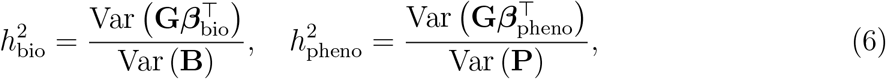

where 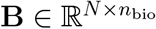 and 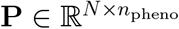 are the matrices of biomarker values and phenotypic values, respectively.

The genetic correlation among biomarkers (***ρ***_**B**_), among phenotypes (***ρ***_**P**_), and between biomarkers and phenotypes (***ρ***_**BP**_) for shared genetic variants were calculated as:

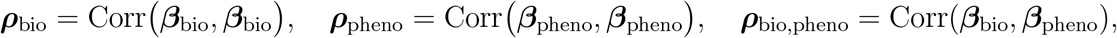

where, for example, the correlation is calculated as:

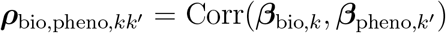

#### 2.2.4 Association testing for simulated traits

Genome-wide association testing for simulated traits was conducted using per-variant linear regression analyses. Specifically, the *k* th biomarker ***b***_*k*_ or phenotype ***p***_*k*_ was associated with the *j*th column of the genotype matrix ***G***_*j*_ according to the model:

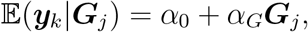

where ***y***_*k*_ *∈* ℝ^*N×*1^ is either ***b***_*k*_ or ***p***_*k*_, *α*_0_ is an intercept, *α*_*G*_ is the genetic effect. For each ***G***_*j*_, representing a single genetic variant, the null hypothesis *H*_0_ : *α*_*G*_ = 0 was evaluated via a standard Wald test. Genome-wide significance was d eclared a t the Bonferroni threshold of 0.05*/J*, where *J* is the total number of variants tested for association.

Across a simulated GWAS, the true positive rate (TPR) was defined as the ratio of the number of causal variants that reached genome-wide significance to the total number of causal variants, The false discovery (FDR) was defined as ratio of the number of non-causal variants that reached genome-wide significance to the total number of variants that reach genome-wide significance.

### 2.3 Methods for real data experiments

#### 2.3.1 Genetic data processing and analyses

To avoid confounding due to population structure, the UK Biobank (UKB) was subset to unrelated subjects of White-British ancestry [19, 18]. Imputed genotypes were filtered to those having a minor allele frequency *>* 1%, INFO score *>* 0.8, and Hardy-Weinberg equilibrium *P >* 1 *×* 10^*−*10^. The following standard covariates were included in all GWAS: age, sex, genotyping array, and the top 20 genetic principal components [18]. Genome-wide association studies (GWAS) and clumping were performed using `PLINK` (v1.9) [20]. GWAS for quantitative traits were performed with linear regression models, and GWAS for binary traits with logistic regression models. Genetic correlation between two traits was estimated using `LDSC v1.0.1` with default settings [21].

#### 2.3.2 Phenotype preparation

Height (UKB: 50), weight (UKB: 21002), and circulating urate (UKB: 30880) were obtained directly from the UKB, filtered non-missing values, and rank-normal transformed [22]. As algorithmically-defined gout was not directly available from UKB, a gout phenotype was constructed following [23]. Specifically, a patient was labeled as having gout if they satisfied at least one of: (1) self-reported gout (code: 1466; UKB: 20002), (2) had an ICD10 code for gout (code: M10; UKB: 41202, 41204, 41270), (3) reported taking allopurinol (1140875408), sulfinpyrazone (1140909890), or colchicine (1140875486) in field 20003, and (4) did not have a hospital diagnosis of leukaemia or lymphoma (codes: C81–C96).

#### 2.3.3 Generating proxy phenotypes with specified target-phenotype correlation

A noisy phenotype *Y*_*ρ*_ having specified correlation *ρ* with a target phenotype *Y* can be generated via:

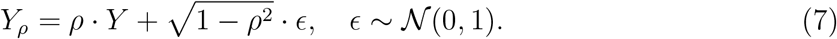

Here *ρ* ∈ [0, 1] controls the expected correlation between *Y*_*ϵ*_ and *Y*, and *ϵ* is mean-zero noise generated independently of *Y*.

#### 2.3.4 Generating a purely environmental proxy phenotype

To construct an imputed phenotype with minimal autosomal heritability, a linear model was fit to predict circulating creatinine (UKB: 30700), a byproduct of protein catabolism, on the basis of age (UKB: 21022), genetic sex (UKB: 22001), and daily alcohol intake, computed as in [24]. The model inputs included polynomial features up to 2nd degree, including all quadratic terms and pairwise interactions:

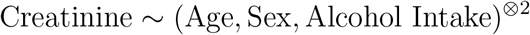

To modulate their association with measured creatinine *Y*, the predicted creatinine levels *Ŷ* were standardized to have mean 0 and unit variance, then corrupted by introducing noise as in (7).

## 3 Results

### 3.1 Downstream vs. Upstream Biomarkers for imputation - which to use?

We first explored how the causal relationship of biomarkers to the true phenotype affects genetic discovery when a machine-learned proxy phenotype, imputed from biomarkers, is studied in place of the true phenotype. **Figure 2** depicts the causal diagrams for the different data generating processes studied, along with the empirical true positive rate (TPR) and the false discovery rate (FDR) as a function of the heritability of the true phenotype *P* and the mean heritability of the biomarkers. We consider four scenarios, in each case using GWAS on the proxy phenotype *P*^*∗*^ as a means of identifying genetic variants associated with the true phenotype *P*. In scenarios (A) and (B), the biomarkers *B* utilized to generate the proxy phenotype lie upstream of the true phenotype on the causal pathway. In (A), the effect of genotype on the true phenotype is fully mediated by the biomarkers, whereas in (B) genotype affects *P* both directly and indirectly via *B*. In both (A) and (B), the FDR is zero. This is because all variants causal for the biomarkers are in fact causal for the true phenotype when *B* lies upstream *P* on the causal pathway. However, scenarios (A) and (B) differ with respect to the TPR. When the effect of genotype is fully mediated by biomarkers as in (A), all variants causal for true phenotype are also causal for the biomarkers. Thus, by studying a proxy phenotype *P*^*∗*^ that is a composite of *B*, it should be possible to recover all variants causal for *P*. As the sample size and biomarker heritability increase, the TPR in scenario (A) will approach 1. In contrast, when the effect of genotype is only partially mediated by the biomarkers as in (B), there exist variants with effects on *P* that do not have effects on *B*. Even with increasing sample size and biomarker heritability, it is not expected that variants whose effects on *P* are not mediated by *B* can be detected by studying *P*^*∗*^. In general, the TPR in (B) will be bounded above by the fraction of variants causal for the true phenotype whose effects are mediated by the biomarkers included in the proxy phenotype.

**Figure 2:**
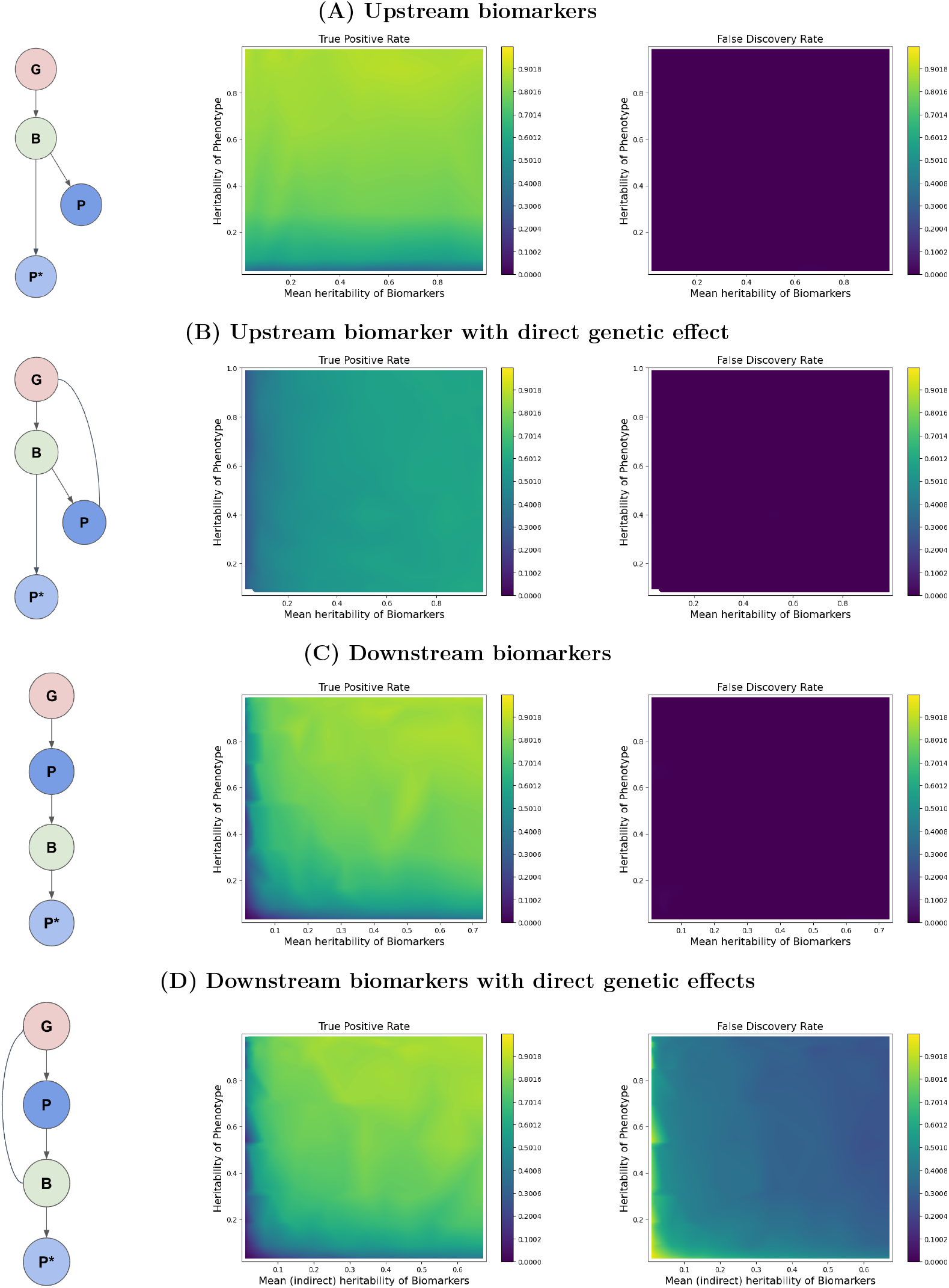
The operating characteristics of proxy GWAS depend on the causal relationship between the true and proxy phenotypes. Four different causal scenarios are show. In (A) and (B), the biomarkers *B* causally upstream of the true phenotype *P*, while in (C) and (D) the biomarkers are causally downstream. In all cases, the proxy phenotype *P*^*∗*^ is derived from the biomarkers. The edges in the causal diagrams on the left show direct effects. The true positive rate in the central column is the proportion of variants causal for *P* that are detected by GWAS for *P*^*∗*^. The false discovery rate in the right column is the proportion of genome-wide significant variants for *P*^*∗*^ that are not causal for *P*.

In scenarios (C) and (D), the biomarkers *B* utilized to generate the proxy phenotype lie downstream of the true phenotype on the causal pathway. In (C), the effect of genotype on the biomarkers is fully mediated by the true phenotype, while in (D) *G* affects *B* both directly and via *P*. In both (C) and (D) the TPR is high, and will approach 1 as the sample size and biomarker heritability increase. This is because all variants causal for the true phenotype are ultimately causal for the collection of biomarkers. Thus, by studying a proxy phenotype derived from the downstream biomarkers, all variants causal for *P* can be detected. Where (C) and (D) differ is with respect to the FDR. In (C), there are no variants with effects on *B* that do not have effects on *P*. Consequently, the FDR is again zero. However, when the effect of genotype on the biomarkers is only partially mediated by the true phenotype as in (D), there exist variants with effects on *B* that do not have effects on *P*. Such variants, which are not relevant to *P*, are expected to surface in GWAS of *P*^*∗*^ as sample size and biomarker heritability increase. In general, the FDR in (D) will be bounded below by the fraction of variants causal for the biomarkers included in the proxy phenotype whose effects are not mediated by the true phenotype.

To illustrate the ideas examined by these simulations, we considered GWAS of two pairs of traits whose causal relationship is well established: → Urate Gout and Height → Weight. For clarity, we consider the simplest possible case, where a single trait directly serves as the proxy for another. GWAS were conducted among 350K unrelated subjects in the UK Biobank, and results were clumped to identify independent (*R*^2^ ≤ 0.1) genome-wide significant (GWS; *P <* 5 *×* 10^*−*8^) loci. Loci for the target trait are considered true positives while loci for the proxy trait are considered predicted positives. Overlap analysis was performed to determine how many loci for the target trait were overlapped by a locus for the proxy trait, and vice versa, from which the empirical TPR and FDR were calculated.

In the case of urate and gout, there were 956 loci for urate and 47 loci for gout. The small number of loci for the latter is attributable to gout being binary rather than continuous, and having low prevalence in our cohort (7.5K cases, 1.5% of the cohort). Viewing urate as the proxy phenotype and gout as the true phenotype, all 47 loci for gout were overlapped by a locus for urate, giving an empirical TPR of 100% (47/47 loci). Of the 956 loci for urate, 590 (61.7%) did not overlap with a locus for gout. There are two possible explanations for the appearance of nominal false positives in the setting of an upstream proxy phenotype. If the measured circulating urate phenotype is truly causal for gout, then the 590 loci detected for urate but not gout are in fact causal for gout, and the latter GWAS simply lacked power to detect them. Alternatively, the measured circulating urate phenotype may be insufficiently specific, for example because only the concentration of urate in the synovial fluid contributes to gout pathogenesis. Causally, this would correspond to a diagram like that in **Figure S1**, where *B*_1_ (e.g. synovial urate) is the biomarker directly on the causal path for gout, and *B*_2_ (e.g. circulating urate) is a genetically correlated biomarker not on the causal path. Studying a proxy phenotype based on *B*_2_ would still allow for detection of true positives *G*_12_ affecting both *B*_1_ and *B*_2_, but will miss true positives *G*_1_ affecting *B*_1_ and not *B*_2_ while introducing the possibility of detecting false positive variants *G*_2_ that affect *B*_2_ and not *B*_1_.

In the case of height and weight, there were 4100 loci for height and 1118 loci for weight. To complement the analysis of urate and gout, we consider weight as a downstream proxy for height. Among the 4100 loci for height, 2308 were overlapped by a locus for weight, for an empirical TPR of 56.3%. Meanwhile, among the 1118 loci for weight, 321 did not overlap a locus for height, for an empirical FDR of 28.7%. Since any locus that affects height should have an effect on weight, failure to detect some height loci via weight is likely due to lack of power, perhaps because more of the variation in weight is attributable to the environment. Consistent with this hypothesis, the heritability of height, as estimated by LD score regression applied to our summary statistics, was 41.5%, compared with only 24.2% for weight. Conversely, those loci for weight that do not overlap a locus for height may affect weight via a pathway other than height, for instance by changing body composition.

### 3.2 Recovery of true positive variants depends on target phenotype heritability and proxy phenotype correlation

We next examine how the TPR depends on the quality of the proxy phenotype, as quantified by its phenotypic correlation with the true phenotype. For the simulation study, we focus on a data generating process in **Figure 3A**, where the proxy phenotype *P*^*∗*^ is imputed from a biomarker *B* purely downstream of the true phenotype *P*. Other data generating processes lead to qualitatively similar conclusions. In the simulation, there are two sources of environmental variation, *E*_1_ affecting *P*, and *E*_2_ affecting *B*. We examine how the TPR depends on the heritability of *P* and the correlation between *P* and *P*^*∗*^. This correlation is in turn determined by the relative influence of *P* versus *E*_2_ on the biomarker that enters the imputation model. For simplicity, we let *B* itself act as the proxy phenotype. The results, presented in **Figure 3B**, demonstrate that success in recovering the causal variants increases with the heritability of the true phenotype and with the correlation of *P* with *P*^*∗*^. Intuitively, the TPR increases as the variation in the proxy phenotype explained by the variants *G* causal for the true phenotype *P* increases. The TPR decreases as more of the variation in *P*^*∗*^ is explained by the environment, either due to having a noisier phenotype (*E*_1_) or due to having a noisier biomarker (*E*_2_).

**Figure 3:**
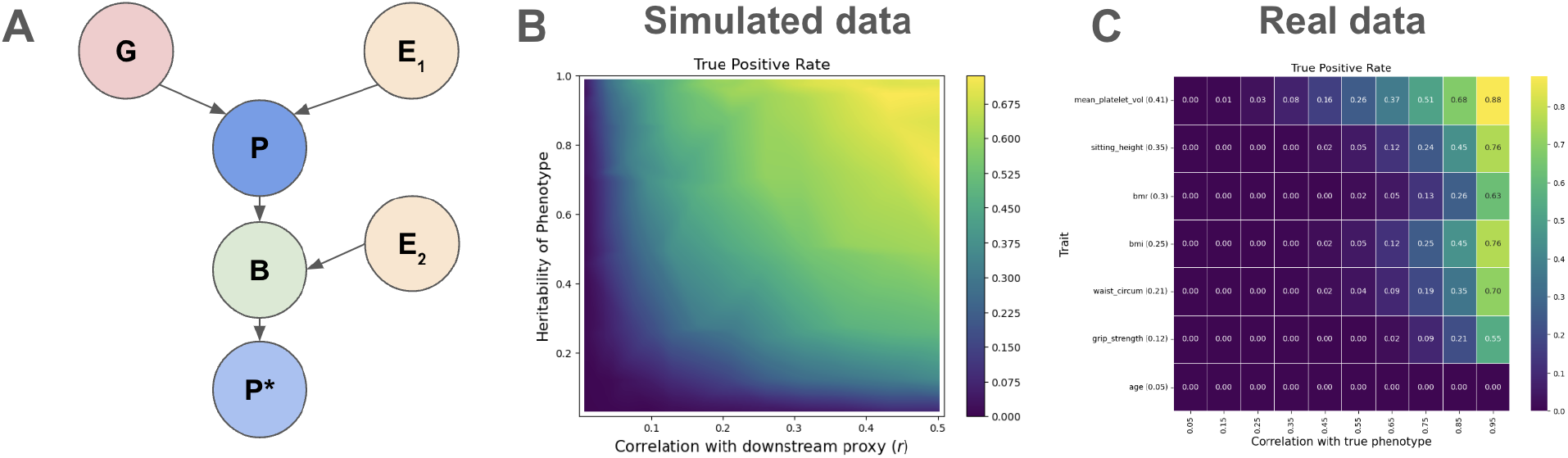
True positive rate increases with target phenotype heritability and proxy phenotype correlation. (A) Causal diagram for the data generating process used in the simulation study. Here *G* is genotype, *P* is the true phenotype, which is affected by environmental factor *E*_1_, *B* is the biomarker, which is affected by environmental factor *E*_2_, and *P*^*∗*^ is the proxy phenotype. (B-C) True positive rate as the proportion of causal variants for *P* recovered as a function of the heritability of *P* and the phenotypic correlation between *P* and *P*^*∗*^. (B) presents results on simulated data, and (C) on real data, where mean-zero noise was introduced to control the correlation between the true and proxy phenotypes.

To illustrate these trends with real data, we selected multiple phenotypes from [25] with heritabilities ranging from 5% for chronological age to 41% for mean platelet volume. Downstream proxy phenotypes having specified correlation with the true phenotype were generated by adding mean-zero noise to the true phenotype, via equation (7), such that the correlation between *P* and *P*^*∗*^ was set to *ρ* ∈ (0.05, 0.10, …, 0.95). The TPR was measured as the proportion of phenotype genome-wide significant for the original *P* recovered via GWAS of *P*^*∗*^. The results, depicted in **Figure 3C**, recapitulate the trends from the simulation study. Recovery of the variants causal for the original phenotype increased with the heritability of *P* and with the correlation between *P* and *P*^*∗*^.

### 3.3 Why phenotypic correlation is a misleading indicator of utility for genetic discovery

The previous experiment suggests that power to detect genetic variants associated with the target phenotype *P* by means of a proxy phenotype *P*^*∗*^ generally increases with the phenotypic correlation between *P* and *P*^*∗*^. However, as we show next, the source of this correlation matters. A proxy phenotype whose correlation with the target is purely environmental rather than genetic in origin will not assist in identifying genetic variants associated with *P*, regardless of the correlation between *P* and *P*^*∗*^. To illustrate this, we simulated a target phenotype *P* and a biomarker *B* according to the data generating process in **Figure 4A**. In contrast to our previous case studies, here the biomarker is neither directly upstream nor downstream of the target phenotype. The proportion of variants having causal effects on both *P* and *B* was varied between 0% and 100%, as was the magnitude of the environmental correlation. The heritabilities of both *P* and *B* were fixed at 50%, and the genetic correlation of *P* and *B* across the subset of variants causal for both phenotypes was fixed at *ρ* = 0.5. The biomarker served as the proxy phenotype.

**Figure 4:**
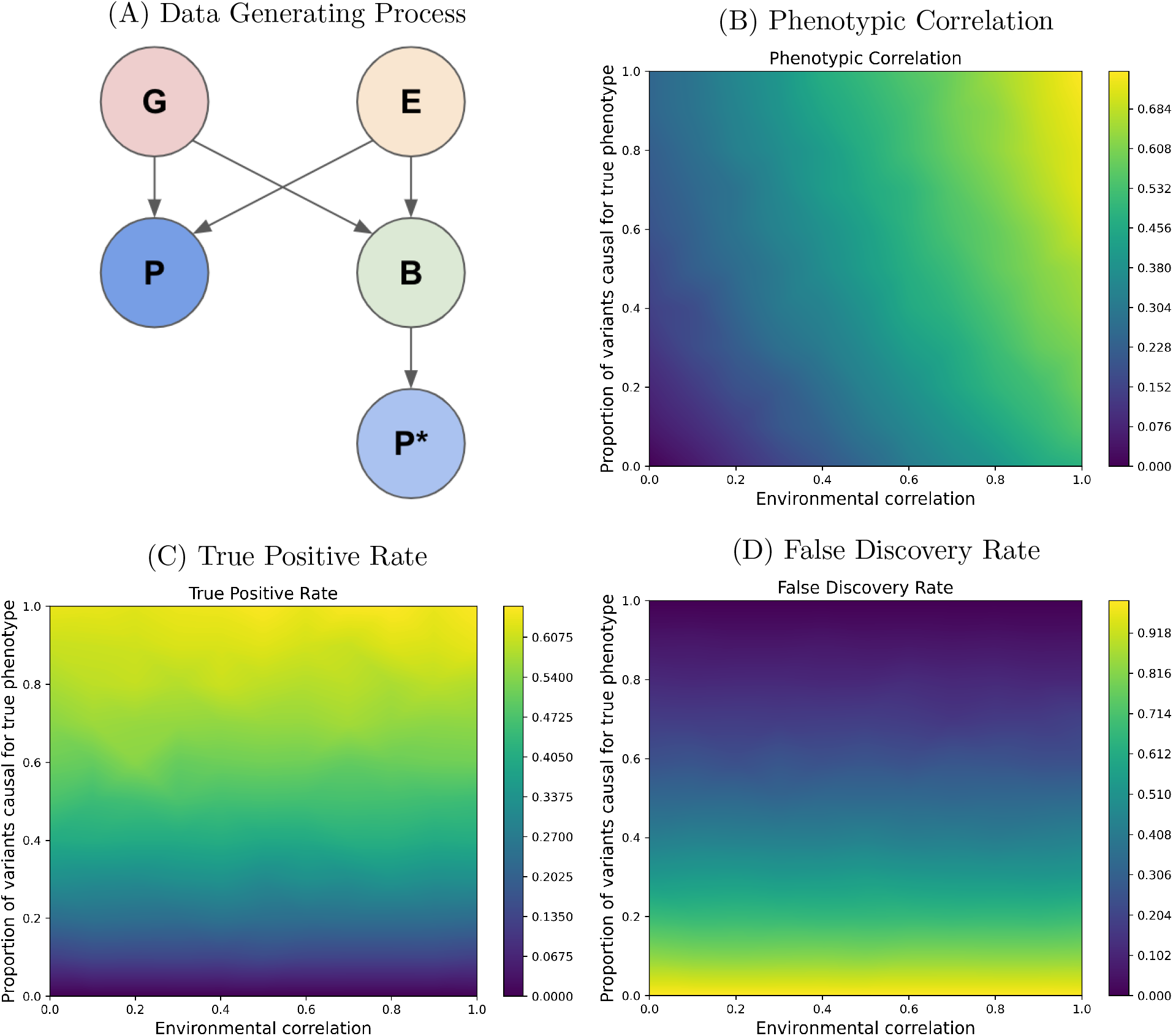
Common genetic basis rather than raw phenotypic correlation determines utility for genetic discovery. (A) Causal diagram for the data generating process. Here *G* is genotype, *E* is the environment, *P* is the target phenotype, *B* is the biomarker, and *P*^*∗*^ is the proxy phenotype. Both *P* and *B* have a heritability of *h*^2^ = 50% and a genetic correlation of zero. The proportion of variants affecting both *P* and *B* and the environmental correlation were each varied between 0 and 100%. (B) Phenotypic correlation as a function of the proportion of shared variants and the environmental correlation. (C) True positive rate and (D) false discovery rate as a function of the proportion of shared variants and the environmental correlation.

**Figure 4B** illustrates how the phenotypic correlation between *P* and *P*^*∗*^ increases with both the proportion of shared causal variants and with the environmental correlation. **Figures 4C** and **4D** demonstrate that increasing the phenotypic correlation by increasing the proportion of shared causal variants increases the TPR and decreases the FDR. Meanwhile, increasing the phenotypic correlation by increasing the environmental correlation has no impact on either the TPR or FDR. Taken together, these results imply that having high phenotypic correlation with the target phenotype is not sufficient for a proxy phenotype to be useful for genetic discovery. In fact, having a high phenotype correlation is also not necessary. Proxy phenotypes with only modest phenotypic correlation but high genetic overlap (i.e. phenotypes in the upper left of **Figure 4B**) will provide greater TPRs and lower FDRs than phenotypes with high phenotypic correlations but poor genetic overlap (i.e. phenotypes in the lower right of **Figure 4B**).

To emphasize the distinction between genetic and environmental correlation, we simulated a phenotype *P*_*G*_ that was predominantly genetic in origin (99% heritability) and a phenotype *P*_*E*_ that was entirely environmental in origin (0% heritability). The data generating processes are shown in **Figure 5A**. A biomarker for *P*_*G*_ was created by adding noise to the genetic component of *P*_*G*_, and a biomarker for *P*_*E*_ was created by adding noise to the environmental component of *P*_*E*_. **Figure 5B** demonstrates that the genetic correlation of the 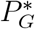 with *P*_*G*_ and of 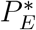 with *P*_*E*_ is stable across a broad range of phenotypic correlations. This means that phenotypic correlation is not an effective substitute for genetic correlation. To illustrate this point with real world data, we constructed a proxy for a highly heritable phenotype by adding noise to a polygenic score (PGS) of height, and a proxy for a minimally heritable phenotype by adding noise to creatinine levels imputed from age, genetic sex, and estimated alcohol intake; creatinine levels are known to be strongly influenced by alcohol intake [26, 27]. **Figure 5C** demonstrates that, as in the simulation study, phenotypic correlation can be toggled independently of genetic correlation. Consequently, we argue that ‘test set *R*^2^’, a common metric for evaluating machine-learning predictions in traditional settings [28], is a poor measure of imputation quality in the context of genetic discovery because it fails to disentangle the genetic signal from the environmental noise.

**Figure 5:**
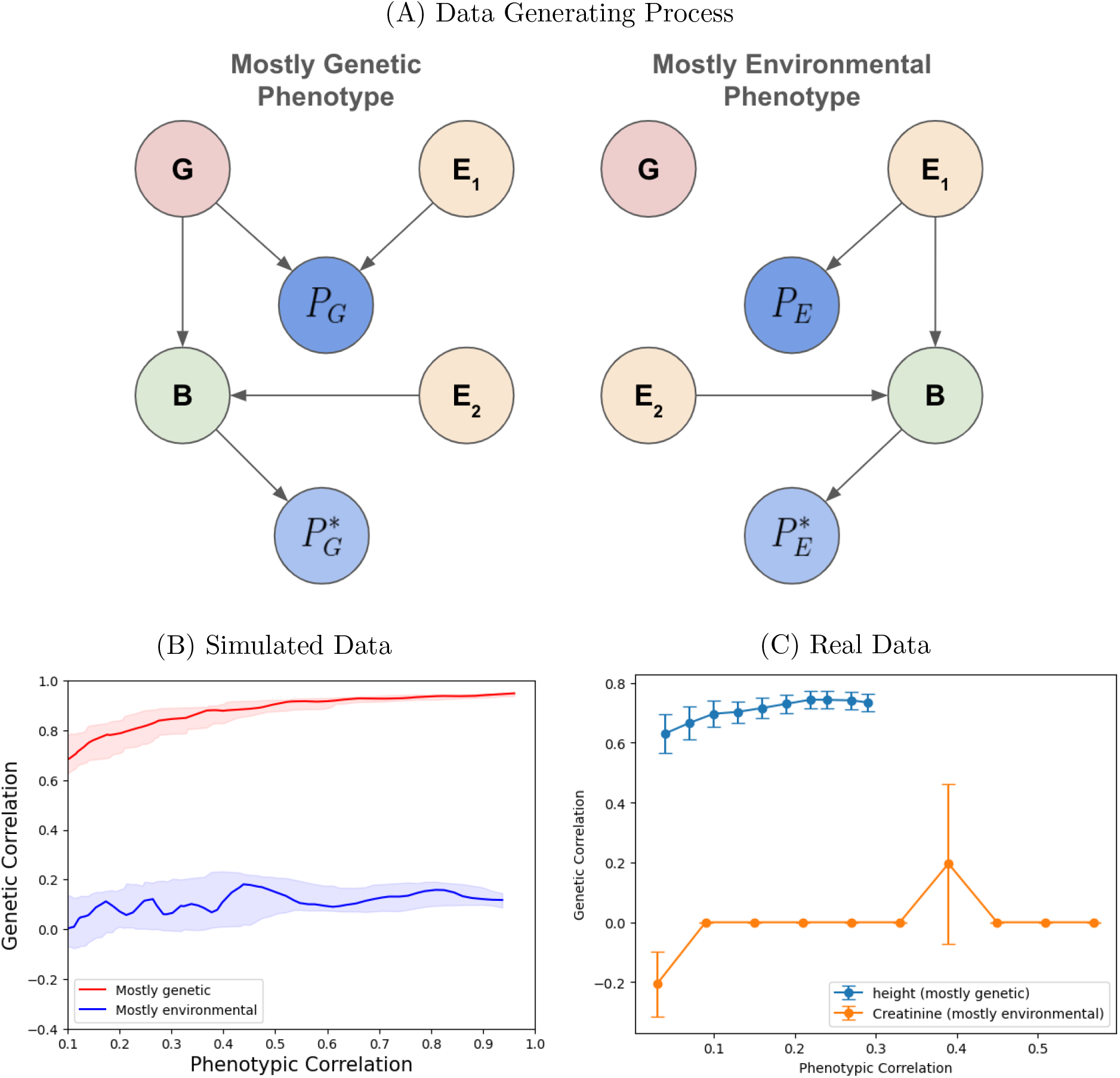
Phenotypic correlation can vary independently of genetic correlation. (A) Causal diagram for the data generating process. Here *G* is genotype, *P*_*G*_ and *P*_*E*_ represent a phenotype that is predominantly genetic or environmental in origin, respectively, 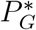 and 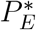 are corresponding proxy phenotypes, *B* is a biomarker, and *E* is environmental noise. (B) and (C) show genetic correlation versus phenotypic correlation for simulated (B) and real (C) data. For the real data setting, a proxy for a highly heritable phenotype was created by adding noise to a polygenic score for height, and a proxy for a minimally heritable phenotype was created by adding noise to circulating createnine imputed from age, genetic sex, and estimated alcohol intake.

## 4 Conclusions

Machine-learning (ML) based imputation presents both opportunities and challenges for genetic discovery. The ability to accurately impute difficult-to-ascertain phenotypes from available surrogate data enables researchers to fill in missing values and augment data sets with unmeasured phenotypes of interest. However, care is needed to ensure the validity of the inferred genetic relationships when conducting genetic association studies with the outputs of ML-models [13, 14, 15]. Here we considered the increasingly common practice of performing GWAS on machine-learned proxy phenotypes. Our analyses of simulated and real data illustrate that considering the causal relationships among genotypes, the phenotype of interest, and any biomarkers input to an ML model is essential to understanding the operating characteristics of proxy GWAS. For example, when the imputation is based on biomarkers known to lie on the causal upstream path of the target phenotype, we can be confident that any associations detected will be relevant to the phenotype of interest. Conversely, when the biomarkers are causally downstream of the target phenotype, proxy GWAS will retain power for detecting true positives but will likely incur contamination by false discoveries. Our work suggests that selecting the appropriate biomarkers for imputation is non-trivial and highly consequential. While using upstream biomarkers for imputation tends to offer better control over false discovery rates, downstream biomarkers can potentially provide higher power for detection, albeit with increased risk of false positives. The strategic choice between these options requires careful consideration of the trade-offs involved and expert knowledge of the underlying pathophysiology of disease. While the true causal relationships between biomarkers and phenotypes may not be always known, large scale mendelian randomization analysis across all biomarkers can provide a rough approximation of what biomarkers might be upstream and which ones might be downstream.

We also demonstrated that although having a higher correlation between the target and proxy phenotypes is generally desirable, the source of the correlation is more important than its magnitude. A proxy phenotype with lower absolute phenotypic correlation that is driven predominantly by genetic overlap will provide greater utility for target-phenotype genetic discovery than a proxy phenotype with higher absolute phenotypic correlation that is driven predominantly by environmental factors. This finding cautions against use of the test set *R*^2^ as a solitary measure of phenotype quality due to its inability to differentiate between genetic and environmental correlations. When evaluating machine-learned proxy phenotypes, having high genetic correlation with the target phenotype is the most desirable scenario, as it suggests the target and proxy phenotypes share many associated variants in common, and that the estimated directions of effect are consistent. Conversely, having a high phenotypic correlation but low genetic correlation is undesirable, as it suggests the correlation is driven by environmental factors, and that many variants associated with the proxy phenotype may not be relevant to the target phenotype. Global genetic correlation, however, is an imperfect indicator of whether a candidate proxy phenotype has high genetic overlap with the target phenotype, as shown by recent work on retinal epithelium pigmentation versus thickness [29]. For the purposes of genetic discovery, the ideal proxy phenotype will associate with all variants that affect the target phenotype, and no variants that do not affect the target phenotype, but need not have the same magnitude or direction of effect. Adding the latter requirements (i.e. same magnitude and direction of effect) would imply a strong (global) genetic correlation with the target and proxy phenotypes.

While this paper has focused on applications of ML in imputing unmeasured or missing target phenotypes, another emerging use of ML in GWAS is to derive lower dimensional representations of high dimensional phenotypes [12, 30, 31, 32, 33, 34]. Due to the difference in the problem specification, careful consideration is likely needed to define what constitute true and false positives for such phenotypes. An important direction of future work is to explore the causal considerations that determine the utility of representation phenotypes for genetic discovery.

## Acknowledgements

The authors would like thank the participants of the UK Biobank, whose data were used with permission. This research was conducted using the UK Biobank Resource under approved Application Number 51766.

