## Supplementary methods for "Causal considerations can determine the utility of machine learning assisted GWAS"

### Supplemental Figures

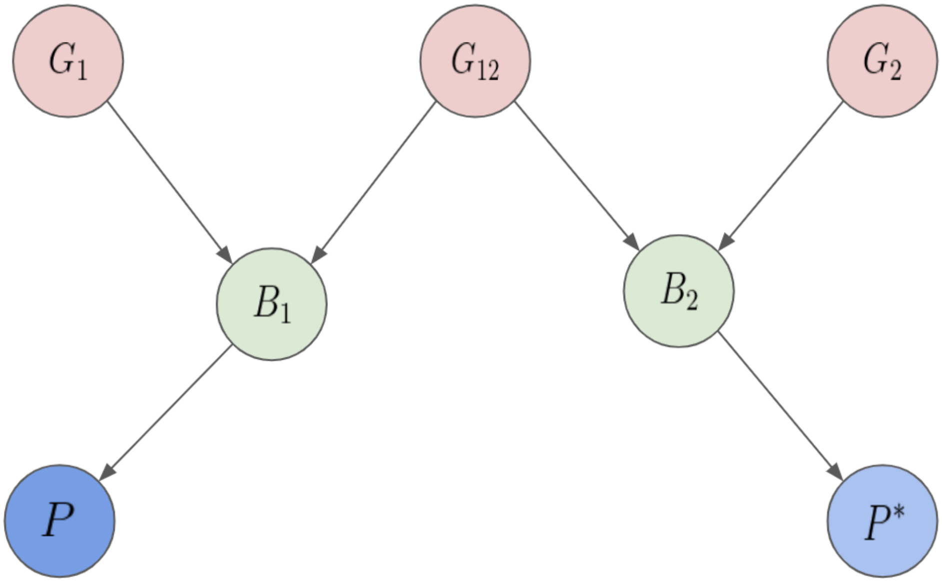


Figure S1: **Causal diagram for an upstream non-specific biomarker.** $B_1$ represents the biomarker (e.g. synovial urate) truly causal for the target phenotype $P$ (e.g. gout) and $B_2$ a related but not causal biomarker (e.g. circulating urate). Variants $G_{12}$ affect both $B_1$ and $B_2$ while variants $G_1$ are specific to $B_1$ and variants $G_2$ to $B_2$. A proxy phenotype $P^*$ based on $B_2$ will provide power to detect variants $G_{12}$ affecting both $B_1$ and $B_2$, but will lack power to detect some true positives $G_1$, and will introduce false positives $G_2$.
